# Association of gut microbiota with metabolism in juvenile Atlantic Salmon

**DOI:** 10.1101/2020.02.10.941245

**Authors:** H. Dvergedal, S.R. Sandve, I.L. Angell, G. Klemetsdal, K. Rudi

**Affiliations:** Department of Animal and Aquacultural Sciences, Faculty of Biosciences, Norwegian University of Life Sciences, P. O. Box 5003, NO-1433, Aas, Norway; Faculty of Chemistry, Biotechnology and Food Science, Norwegian University of Life Sciences, P. O. Box 5003, NO-1433, Aas, Norway

**Keywords:** microbiome, Atlantic salmon, genetics, metabolism, feed efficiency, carbon turnover

## Abstract

The gut microbiome plays a key role in animal health and metabolism through the intricate functional interconnection between the feed, gut microbes, and the host. Unfortunately, in aquaculture, the links between gut microbes and fish genetics and production phenotypes are not well understood.

In this study, we investigate the associations between gut microbial communities, fish feed conversion, and fish genetics in the domestic Atlantic salmon. Microbial community composition was determined for 230 juvenile fish from 23 full-sib families and was then regressed on growth, carbon and nitrogen metabolism, and feed efficiency. We only found weak associations between host genetics and microbial composition. However, we did identify significant (*p* < 0.05) associations between the abundance of three microbial operational taxonomical units (OTUs) and fish metabolism phenotypes. Two OTUs were associated with both carbon metabolism in adipose tissue and feed efficiency, while a third OTU was associated with weight gain.

In conclusion, this study demonstrates an intriguing association between host lipid metabolism and the gut microbiota composition in Atlantic salmon.

## Background

Efficient and environmentally sustainable animal production systems are urgently required to ensure long-term food security, especially as global aquaculture consumption is projected to double by 2050 (www.fao.org). One important aspect of improving sustainability is to improve feed conversion and growth. In humans and other vertebrate systems, the gut microbiome plays a central role in the path from “feed-to-animal” (1–4), and recent studies have also shown that host-genetic factors can modulate microbiome composition. Such functional interconnection between feed, microbes, and host (i.e. the feed-microbiome-host axis), opens up intriguing avenues for optimizing aquaculture production systems, for example by breeding for ‘optimized’ microbiome composition (5).

Yet, even though the dietary composition is known to impact the gut microbiome in aquaculture species (1), almost nothing is known about the link between the gut microbiota and important production phenotypes, or to what extent microbiota composition itself could be a new breeding target for aquaculture breeding programs (5).

To address this pressing knowledge gap we use a family-based experimental design to test if variation in the gut microbiome composition in juvenile Atlantic salmon is associated with key phenotypes related to host metabolism as well as variation in host genetics. Our results identified phenotypic associations between host gut microbiome and lipid metabolism, growth, as well as to feed efficiency, which open the possibility for metabolic modulation through the gut microbiota.

## Materials and methods

### Experimental setup

A family experiment with Atlantic salmon was carried out at the fish laboratory, Norwegian University of Life Sciences (NMBU), Aas, Norway, according to the laws and regulations controlling experiments on live animals in EU (Directive 2010/637EU) and Norway (FOR-2015-06-18-761). The experiment was approved by the Norwegian Food Safety Authority (FOTS ID 11676).

The family experiment is explained in detail by Dvergedal et al (6). In short, broodstock from AquaGen’s breeding population (22 males and 23 females) were used to generate 23 families. To ensure clearly contrasted family groups with respect to growth potential, the parents were selected in two directions for high and low estimated breeding values (EBVs) for growth in seawater, respectively.

Prior to the start-feeding several families were kept in separate compartments within the same tank, and five tanks were needed to house all families. Based on parentage assignment 100 family members were identified for each of the 23 families and reared together in a single tank from start-feeding until the start of the experiment. *A priori* to the 12-day test, families were allocated to tanks, 50 fish per tank and 2 tanks per family (except for nine tanks in which the number of fish varied between 42 and 54, due to some mortality prior to the start of the experiment or an increased number due to a counting mistake). From each tank five fish (10 fish per family) were collected for microbiota and phenotypic analyses, a total of 230 fish were sampled all together (Fig. 1). Families were fed a fishmeal-based diet labeled with the stable isotopes ^15^N and ^13^C, with inclusion levels of 2% and 1%, respectively, as described in Dvergedal et al. (6).

**Figure 1.**
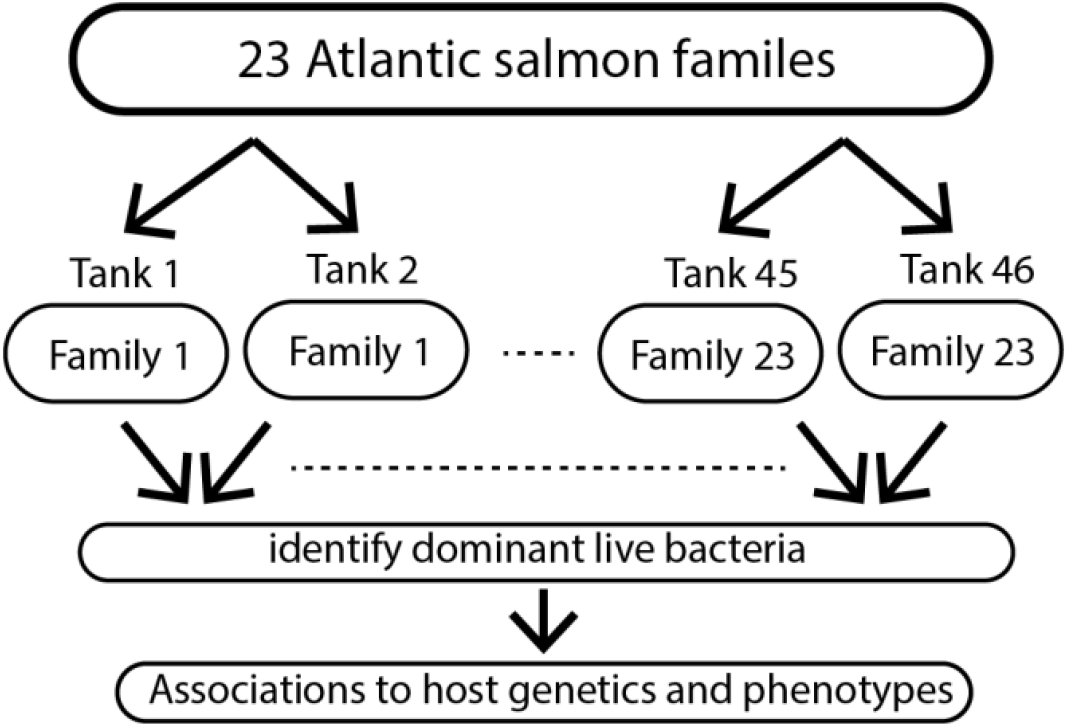
Schematic outline of the experimental setup. Twenty-three families were distributed on 46 tanks (two tanks per family). Dominant live bacteria were identified prior to association analyses to fish genetics and metabolism.

The tanks, each with a 270-L capacity, were supplied with water from a common source of recirculated fresh water, at a flow rate of 7 to 8 L.min^-1^. The fish were kept under 24 h light regime, with an average temperature of 14.5°C. Dissolved oxygen was measured daily and maintained above 8 mg.L^-1^ in the outlet water (Handy Delta, OxyGuard^®^ AS, Farum, Denmark).

### Microbiota analyses

Distal intestinal samples (n = 230) were obtained by squeezing out the gut segment content using sterile tweezers in 1 ml phosphate buffered saline (PBS) and put on ice until further processing. To distinguish between DNA from dead and alive bacteria, the samples were treated with propidium monoazide (PMA) within eight hours post sampling in order to inactivate free DNA, and DNA in dead cells (7). Samples were pulse centrifuged up to 1200 rpm and split in two, where one part (n = 230, PMA treated samples) was added PMA dye (Biotium, USA to a final concentration of 50 µM, and the other part was kept as a control (n = 230, non-PMA treated samples) with no added PMA. The samples were then kept dark for 5 min before exposure to light for 30 min in a lightbox from Geniul. DNA extraction (n = 460) was done using mag midi DNA extraction kit (LGC Genomics, UK) following the manufacturer’s recommendations.

The 16S rRNA amplicon library was prepared and sequenced as previously described (8). Briefly, this involved amplification in 25-µl volumes, with 0.2 µM of both primers, and 2 µl genomic DNA. The PCR cycles involved denaturation at 95°C for 30 s, annealing at 55°C for 30 s, with an initial heat activation at 95°C for 15 min. Illumina modified adapters added with 10 new PCR cyles after purification with AMPure XP beads (Beckman-Coulter, USA) were. Negative controls without genomic DNA were included on all PCR plates (n=5), and included in sequencing if giving detectable band by agarose gel electrophoresis. The sequence reads were processed using USEARCH v8 (9) where the sequences were paired-end joined, demultiplexed, and quality filtered (maxxee = 1.0, minlength = 350, singletons discarded), before operational taxonomic unit (OTU) clustering with 97% identity threshold was performed using the UPARSE pipeline (10). Taxonomy assignment was done using SILVA database (11). Diversity analysis was done using a sequence depth of 10 000 sequences per sample. These analyses were done using default parameters.

To filter out OTUs from dead bacteria and bacteria considered as contaminants, filtering was done using the following criteria on each individual fish gut microbiome; OTUs which showed a more than 3-fold reduction in the PMA treated sample was considered dead, while OTUs that showed a more than 6-fold increase in the non PMA treated sample were considered contaminants because there were no other alternative explanations. Out of the 230 fish gut microbiomes, 188 passed the sequence quality control filtering criteria, including rarefaction at 10 000 sequences as a tradeoff between number of samples and sequencing dept, in addition to live/dead/contamination screening. Correlations between OTUs were determined using Spearman’s rank correlation coefficient. Raw16S rRNA sequence data are deposited in the SRA database under the accession number PRJNA590084.

### Phenotypic data

The host metabolism related traits analyzed are listed in Table 1. Details for phenotypic data for growth and metabolic traits are explained in Dvergedal et al. (6).

**Table 1.** Description of the 13 variables phenotyped.

| No. | Variables | Description |
| --- | --- | --- |
| 1 | IW | Initial weight <sup>2</sup> |
| 2 | FW | Final weight <sup>2</sup> |
| 3 | WG | Weight gain <sup>2</sup> (FW-IW) |
| 4 | RG | Relative weight gain (%) (((FW-IW)/FW)100) |
| 5 | AMC | Atom % <sup>13</sup>C in muscle |
| 6 | AMN | Atom % <sup>15</sup>N in muscle |
| 7 | ALC | Atom % <sup>13</sup>C in liver |
| 8 | ALN | Atom % <sup>15</sup>N in liver |
| 9 | AAC | Atom % <sup>13</sup>C in adipose tissue |
| 10 | IFCR_AMC | Individual isotope-based indicator of feed conversion ratio, from Atom % <sup>13</sup>C in muscle |
| 11 | IFCR_AMN | Individual isotope-based indicator of feed conversion ratio, from Atom % <sup>15</sup>N in muscle |
| 12 | IFER_AMC | Individual isotope-based indicator of feed efficiency ratio, from Atom % <sup>13</sup>C in muscle |
| 13 | IFER_AMN | Individual isotope-based indicator, of feed efficiency ratio, from Atom % <sup>15</sup>N in muscle |
| 14 | OTU1 | Operational taxonomic unit 1, classified as *Caulobacteriaceae* |
| 15 | OTU2 | Operational taxonomic unit 2 classified as *Pseudomonas fluorescens* |
| 16 | OTU3 | Operational taxonomic unit 3 classified as *Sphingomonas* |
| 17 | OTU5 | Operational taxonomic unit 5 classified as *Bradyrhizobium* |
| 18 | OTU6 | Operational taxonomic unit 6 classified as *Ralstonia* sp |
| 19 | OTU7 | Operational taxonomic unit 7 classified as *Pseudoalteromonas* |

### Outlier detection

To obtain approximate normality of the relative abundances of OTUs we transformed the OTU data using the natural logarithm (Ln). Influence statistic was used for outlier detection by regressing Ln (OTU) on all the phenotypes (Table 1) using PROC REG in SAS^®^. The cutoff value for outliers was calculated as 3*p*/*n* (>0.10), where *n* is the number of samples (i.e. animals) used to fit the model (n = 188), and *p* is the number of parameters in the model. A total of 16 outliers were detected and deleted.

### Estimation of heritability

To estimate heritabilities of the microbiota at the level of each OTU we first did a single-trait analysis of variance of the Ln (OTUs). In each analysis, the model was:

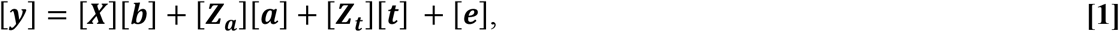

where [***y***] is a vector of individual OUT ‘phenotypes’ (i.e. the trait), [***b***] is a vector of fixed effects, including sampling *day*_*i*_ (*i* = 1-4), 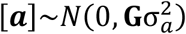 is a vector of random additive genetic effects for the trait, 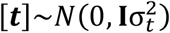 is a vector of random tank effects for the trait, and 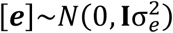, is a vector of random residuals for the trait. The **X** and **Z** matrices are corresponding incidence matrices, 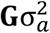 is the genomic (co)variance matrix, 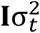 is the (co)variance matrix due to tank effects, and 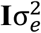 denotes the error (co)variance matrix. The number of phenotyped individuals was rather low (n = 172), and the genomic relationship matrix was generated according to VanRaden’s first method (12). The matrix **G** (2282×2282) was calculated based on a subset of 51,543 SNPs of high genotype quality, covering all autosomal chromosomes (AquaGen’s custom Axiom^®^SNP genotyping array from Thermo Fisher Scientific (San Diego, CA, USA) includes 56,177 single-nucleotide polymorphisms).

Heritabilities of the OTUs were estimated as: 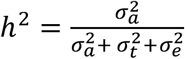, where 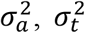, and 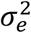 are the estimates of the individual additive genetic, tank environmental, and individual residual variance, respectively, of the trait. The fraction of variance explained by the tank was estimated as: 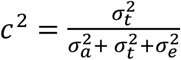. Significance of the genetic effect was tested using a likelihood-ratio (*LR*) test-statistic, comparing a single-trait model with genetic effects (H1) to a model without genetic effects (H0):

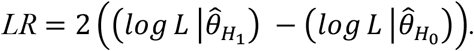

The genetic effect was considered significant if 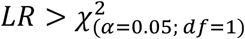.

### Genome-wide association analysis

To associate variation in microbial composition with host genetics a genome-wide association study was done using Ln (OTUs) as response variables. The analysis was carried out by a linear mixed-model algorithm implemented in a genome-wide complex trait analysis (GCTA)^12^. The leave one chromosome out option (*--mlm-loco*) was used, meaning that the chromosome harboring the SNP tested for was left out when building the genetic relationship matrix (GRM). The linear mixed model can be written:

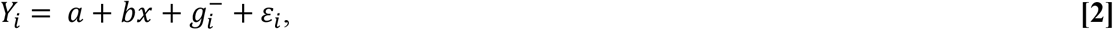

where *Y*_*i*_ is one of the Ln (OTUs) of individual *i, a* is the intercept, *b* is the fixed regression of the candidate SNP to be tested for association, *x* is the SNP genotype indicator variable coded as 0, 1 or 2, 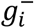 is the random polygenic effect for individual 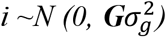 where **G** is the GRM and 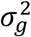 is the variance component for the polygenic effect, and ε*i* is the random residual. In this algorithm, 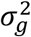 is re-estimated each time a chromosome is left out from the calculation of the GRM. The dataset was filtered, and individuals with < 10% missing genotypes were kept (*n =* 2279). Further, it was required that SNPs should have minor allele frequency (MAF) ≥ 1% and a call rate > 90%. After filtering 54,200 SNPs could be included in the analysis. The level of significance for SNP was evaluated with a built-in likelihood-ratio test, and the threshold value for genome-wide significance was calculated by the use of Bonferroni correction (0.05/54200) = 9.23 × 10^−7^, corresponding to a -log_10_ *p-value* (*p*) of 6.03.

### Association between OTUs and fish phenotypes

We examined the association between microbiota and several individual fish phenotypes related to the metabolism of the fish, including growth, nutrient turnover, and feed efficiency parameters (Table 1). This phenotypic association were tested with a linear mixed-effect model:

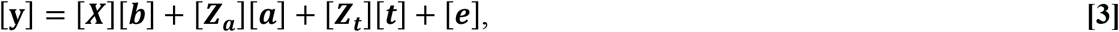

where, [*b*] is a vector of fixed effects for the trait containing the regressions of sampling day (Day) and the Ln (OTUs), while the remaining are described with **[1]**. In preceding analyses, we experienced a strong co-linearity between the tank and the family effects, which in consequence led us to analyze for the phenotypic association between production variables and the OTUs with a model accounting for the genomic relationships between individuals (also known as a genomic selection model).

The genetic analyses in **[1]** and **[3]** were carried out using the ASReml4 software package (14).

## Results

### Overall microbiota composition

We obtained a total of 9 991 266 read counts after filtering and paired-end sequence merging. The mean read count per sample was 22 007. A total of 704 OTUs were defined, with the dead bacterial fraction representing 9.1% of the sequencing reads (contained in 254 OTUs). In addition, a fraction of 0.003% of the sequencing reads (contained in 146 OTUs) were considered as contaminations (Fig. 2A). Among the 304 OTUs passing the live/dead/contamination filtering we identified a clear over-representation of *Proteobacteria*, both with respect to OTU prevalence and quantity (Fig. 2B). OTU1 *Caulobacteraceae* and OTU2 *Pseudomonas fluorescens* dominated, with mean abundances of 32.9% and 34.8%, respectively. There were 6 OTUs with a mean abundance > 1 % in all the samples. We detected an overall negative correlation between OTU1 *Caulobacteraceae* to the other OTUs (Fig. 2C). Bacterial DNA was detected in one out of five negative control, with a dominance of Halomonas (43.4%) and Pseudoalteromonas (40.6 %), followed by Bacillus (4.2 %) and Pseudomonas fluorescens (4.1%).

**Figure 2.**
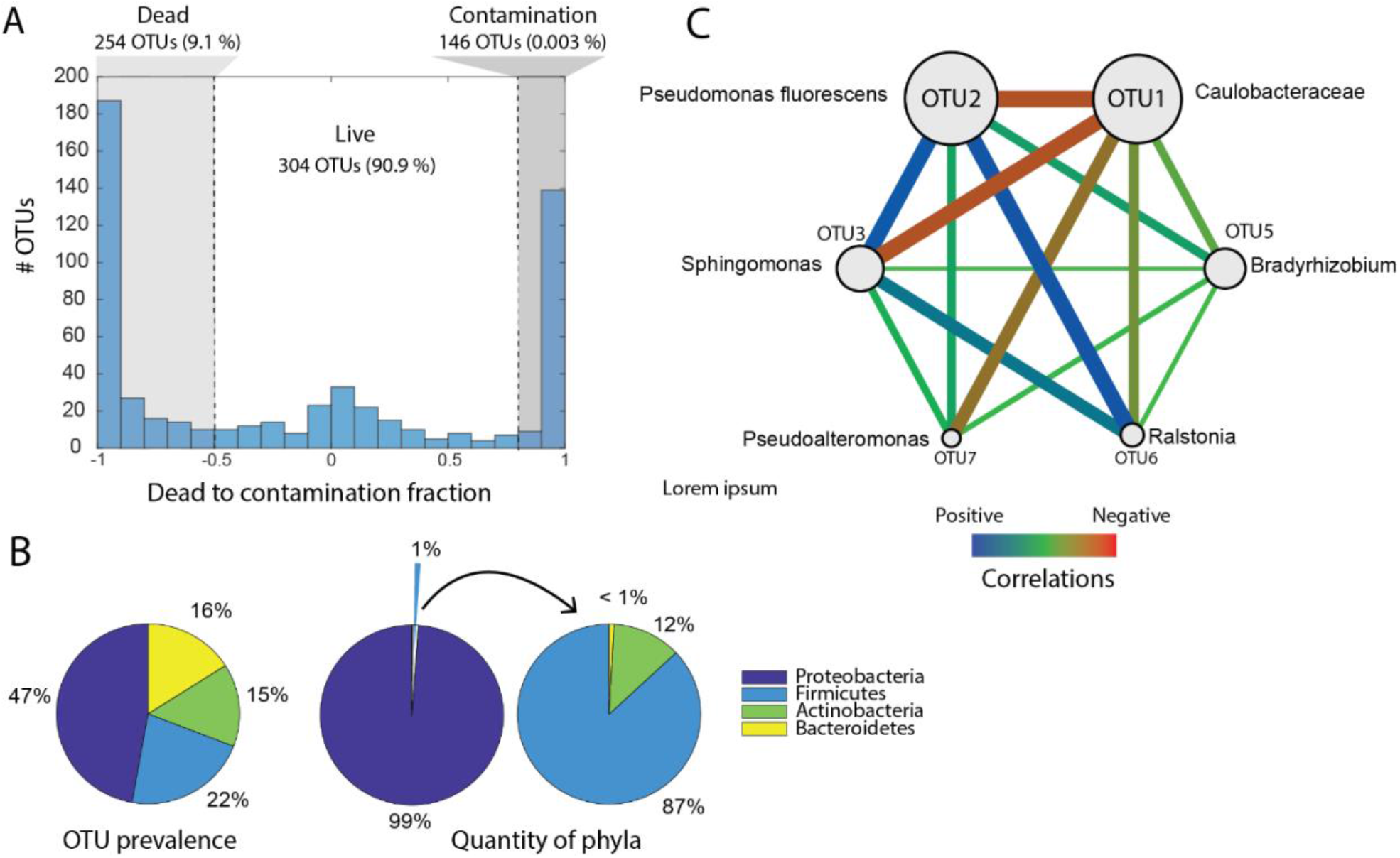
General characteristics of gut microbiota. (A) Fractions of dead, live, and contamination OTUs. Fractions calculated based on the ratio of OTU counts in PMA treated samples versus untreated samples. (B) Prevalence (number of OTUs) and quantity (number of sequencing reads) for the different phyla in the most abundant OTUs. (C) Spearman correlations between the dominant live bacteria (high-abundant OTUs). The high abundant OTUs were identified as those that have an abundance of > 1% on average in all the samples.

### Effects of rearing tank and host genetics on the gut microbiome composition

To assess the contribution of genetics (i.e. heritability) and environment (i.e. tanks effect) in driving the variation in microbial composition between individual fish we applied analyses of variance, using tank as a covariate. Although we classified 304 OTUs across all gut microbiomes, the top 6 most abundant OTUs represented 85% of the 16S sequences in our dataset. We, therefore, conducted these analyses using only these OTUs. The results (Table 2) showed small genetic components for OTUs 1 and 3 and only a small non-significant tank effect for OTU1 (0.03). However, neither the tank nor genetic effects were significantly different from zero (*p* > 0.05). However, the standard errors were large, meaning that the experiment did not have the power to estimate these components precisely. This is supported by the estimates of variance components for genetic or tank effects (or both) of the remaining OTUs being zero (i.e. restricted to the boundary of the parameter space) (Table 2).

**Table 2.** Estimates of the genetic, tank and residual variance components (, and, respectively), the fraction of phenotypic variance explained by the environmental tank effect (), heritability (), as well as the χ^2^ statistics for the additive genetic family effect, with the corresponding level of significance (*p*).

| | $\sigma_a^2$ | $\sigma_t^2$ | $\sigma_e^2$ | $c^2$ | $h^2$ | $\chi^2$ | $p$ |
| --- | --- | --- | --- | --- | --- | --- | --- |
| OTU1 | 0.007 | 0.009 | 0.25 | $0.03 \pm 0.08$ | $0.03 \pm 0.09$ | 0.13 | 0.72 |
| OTU2 | 0.00<sub>1</sub> | 0.00<sub>1</sub> | 0.14 | 0 | 0 | 0 | 1 |
| OTU3 | 0.007 | 0.00<sub>1</sub> | 0.58 | 0 | $0.10 \pm 0.09$ | 1.72 | 0.19 |
| OTU5 | 0.00<sub>1</sub> | 0.00<sub>1</sub> | 0.13 | 0 | 0 | 0 | 1 |
| OTU6 | 0.00<sub>1</sub> | 0.00<sub>1</sub> | 0.07 | 0 | 0 | 0 | 1 |
| OTU7 | 0.00<sub>1</sub> | 0.00<sub>1</sub> | 0.18 | 0 | 0 | 0 | 1 |

<sub>1</sub>Restricted on the boundary of parameter space

Finally, we utilize the existing genotyping data for these fish(6) to perform a genome-wide association analyses for the OTU abundances. No genome-significant associations between SNPs and OTUs were identified, however, the Manhattan plots show clear peaks at chromosomes 14, 24, 3, and 5 (Suppl. Fig. 1), with some SNPs having significant associations to OTU1 and OTU2 at the chromosome level (Suppl. Fig. 2, Suppl. Table 1).

### Community structure is associated with fish growth and metabolism

Linear regressions were used to examine the phenotypic relationship between the gut microbiome and host metabolism traits (i.e. growth, nutrient turnover, and feed efficiency, see table 1). Indeed, these analyses (Table 3) do indicate a link between the production variables and the gut microbiome (significant associations in Table 3, for all results see Suppl. Table 2).

**Table 3.** Regression estimates, standard errors, *F*- and *p*-values when regressing OTUs on growth, metabolism, and feed efficiency variables. The model also contained regression on day and random effects of animal (utilizing genomic relationships) and tank, for which variance components are included.

| Dependent variable | Variables | Estimate | Stderr | F-value | *p*-value | Variance component |
| --- | --- | --- | --- | --- | --- | --- |
| WG | Day | 3.338 | 0.177 | 355.73 | <0.005 |  |
|  | OTU1 | -0.324 | 0.376 | 0.15 | NS |  |
|  | OTU2 | 2.191 | 1.199 | 0.04 | NS |  |
|  | OTU3 | -1.352 | 0.624 | 4.66 | <0.05 |  |
|  | OTU5 | -0.743 | 0.535 | 1.78 | NS |  |
|  | OTU6 | -0.440 | 0.866 | 0.26 | NS |  |
|  | OTU7 | 0.179 | 0.492 | 0.30 | NS |  |
|  | Tank |  |  |  |  | 13.84 ± 3.25 |
|  | Animal |  |  |  |  | 5.54 ± 0.69 |
| AAC | Day | 0.082 | 0.019 | 17.95 | <0.005 |  |
|  | OTU1 | 0.007 | 0.003 | 1.39 | NS |  |
|  | OTU2 | -0.016 | 0.009 | 2.15 | NS |  |
|  | OTU3 | -0.001 | 0.005 | 1.31 | NS |  |
|  | OTU5 | 0.006 | 0.004 | 4.63 | <0.05 |  |
|  | OTU6 | -0.001 | 0.007 | 0.01 | NS |  |
|  | OTU7 | 0.010 | 0.004 | 8.53 | <0.005 |  |
|  | Tank |  |  |  |  | 0.90 ± 0.20 |
|  | Animal |  |  |  |  | 0.87x10<sup>-4</sup> ± 2.21x10<sup>-4</sup> |
| IFER_AMC | Day | 0.365 | 0.021 | 297.49 | <0.005 |  |
|  | OTU1 | 0.043 | 0.026 | 5.55 | <0.03 |  |
|  | OTU2 | -0.101 | 0.083 | 0.41 | NS |  |
|  | OTU3 | 0.018 | 0.043 | 1.34 | NS |  |
|  | OTU5 | -0.023 | 0.037 | 0.03 | NS |  |
|  | OTU6 | 0.035 | 0.060 | 0.34 | NS |  |
|  | OTU7 | 0.081 | 0.034 | 5.22 | <0.03 |  |
|  | Tank |  |  |  |  | 0.20 ± 0.04 |
|  | Animal |  |  |  |  | 0.43x10<sup>-2</sup> ± 1.51x10<sup>-3</sup> |
| IFER_AMN | Day | 0.149 | 0.009 | 273.21 | <0.005 |  |
|  | OTU1 | 0.020 | 0.009 | 3.89 | <0.05 |  |
|  | OTU2 | -0.026 | 0.027 | 2.86 | NS |  |
|  | OTU3 | 0.001 | 0.014 | 0.64 | NS |  |
|  | OTU5 | -0.016 | 0.012 | 0.79 | NS |  |
|  | OTU6 | 0.008 | 0.020 | 0.16 | NS |  |
|  | OTU7 | 0.027 | 0.011 | 5.83 | <0.03 |  |
|  | Tank |  |  |  |  | 0.04 ± 7.86x10<sup>-3</sup> |
|  | Animal |  |  |  |  | 0.29x10<sup>-3</sup> ± 1.51x10<sup>-4</sup> |

OTU3 (*Sphingomonas*) regressed negatively on weight gain (*p* < 0.05), OTU1 (*Caulobacteriaceae*) and OTU7 (*Pseudoalteromonas*) regressed positively on feed efficiency indicators (*p* < 0.05), while OTU7 (*p* < 0.005) and OTU5 (*Bradyrhizobium*) (*p* < 0.05) regressed positively on carbon metabolism in adipose tissue (AAC variable).

## Discussion

A major strength of this experiment is that each individual fish microbiome can be linked to detailed individual-level phenotypes of growth, feed efficiency, and nutrient turnover as measured by the use of stable-isotope profiling in the liver, muscle, and adipose tissues (6). Intriguingly, the phenotypic associations between OTUs and fish production-related phenotypes revealed several significant relationships (Table 3). We observed significant positive associations between lipid carbon metabolism (the AAC phenotype) and OTU5 (*p* < 0.05) and OTU7 (*p* < 0.005). These OTUs belong to the genera *Bradyrhizobium*, and *Pseudoalteromonas*, respectively. *Pseudoalteromonas* is known to have the capacity to produce a range of biologically active extracellular compounds, ranging from antimicrobial compounds and proteases to compounds important for host metamorphosis (15). This genus has also been used as probiotics in fish farming (16). *Bradyrhizobium*, on the other hand, is a widespread environmental bacterium capable of degrading aromatic compounds and nitrogen fixation (17). However, the potential mechanisms for the bacterial associations with lipid metabolism in salmon are completely unknown. The *Sphingomonas* OTU3 showed a significant negative association with weight gain. This genus has previously been associated with antibiotic resistance connected to disease treatment of juvenile salmon in fresh-water (18), which may indicate that reduced weight gain could be connected to the opportunistic properties of *Sphingomonas* (*19*). Lastly, OTU1 belonging to *Caulobacteriacea* showed a strong negative correlation with the other OTU (Fig. 2C), and a significant positive association with two of the feed utilization efficiency metrics (Table 3), which also had a positive association to OTU7. One interpretation of this is that *Caulobacteriacea* has a mutualistic association with juvenile salmon, possibly by protecting juvenile salmon against opportunistic infections.

The association between OTUs and the lipid carbon metabolism (AAC) in Atlantic salmon means that fish with higher abundances of these microbes also convert the feed protein fraction to lipids at a higher rate. However, the mechanisms driving this association remains elusive. One possibility is that the Atlantic salmon gut microbiota has a direct impact on the production of biomolecules in the distal intestine which are readily absorbed and deposited as fat in adipose tissues. It is worth noting that Dvergedal et al. (6) have reported that fish with a higher turnover of carbon in lipid tissues also have improved feed efficiency (see IFER variable in Tables 1 and 3) and fast growth.

In other words, the fish with high carbon turnover in lipid tissues will likely have a positive energy balance and therefore also the opportunity to convert more surplus energy into lipids for storage. We did however not observe associations between gut microbes and nutrient turnover in muscle or liver. This lack of association with nitrogen turnover could be because the majority of the protein fraction is digested and absorbed before the distal intestine where our microbial samples were collected from. It is also possible that the associations between microbial composition and fish metabolism are driven by indirect factors. Since growth is positively correlated with feed intake (6), these fish might also have increased passage rate in the gastrointestinal tract due to high feed intake. This could indirectly affect the competition and balance among microbes and thereby shift the community structure. It is thus critical that future studies include functional meta-omics data that can also demonstrate shifts in activities in microbial metabolic pathways.

Host genetic effects on gut microbiomes have been identified in recent studies in a wide range of animals, including invertebrates (20), mammals (21,22), and fish (23). Here we associate OTUs and metabolic fish phenotypes using a regression, correcting for the effects of additive genotype and tank, aiming to eliminate possible confounding effects. Using this approach, we only found weak (and non-significant) associations between host genetics and relative OTU abundance (Table 2, Supplementary Fig. 1 and 2). Neither did we find rearing tank effects, in correspondence with a study in tilapia (24). However, it is clear from the standard errors (Table 2) that our OTU heritability estimates were very imprecise. Hence, to assess the importance of host genetics on gut microbiome composition in Atlantic salmon, future studies must increase sample size significantly and apply ‘common rearing’ experimental designs to avoid confounding tank and family effects. The PMA screening indicates that more than half of the OTUs detected in the salmon gut could come from feed or dead bacteria. The PMA assay, however, does not cover contaminants in reagents. The negative controls revealed detectable contamination in only one sample, with none of the common reagent contaminants being detected at levels > 1 % (25,26). The most likely source of contamination in that sample would therefore be spillover, and not reagent contaminants.

## Conclusion

In conclusion, our results demonstrate an association between the microbial composition in the distal gut and a key aspect of Atlantic salmon metabolism. This association could be a direct effect of microbes contributing to improved nutrient availability and absorption for the host. Alternatively, these associations could be uncoupled from the microbiota function and instead driven by feeding behavior and passage rates. Future experiments should, therefore, aim to measure changes in microbial metabolic pathways to separate causal from correlative microbe-host associations.

## Acknowledgments

We thank Margareth Øverland, Liv Torunn Mydland and Jørgen Ødegård for the planning of the experiment and Jørgen Ødegård for creating the genomic relationship matrics used in this study.

## SUPPLEMENTARY TABLES

**Supplementary Table 1.**
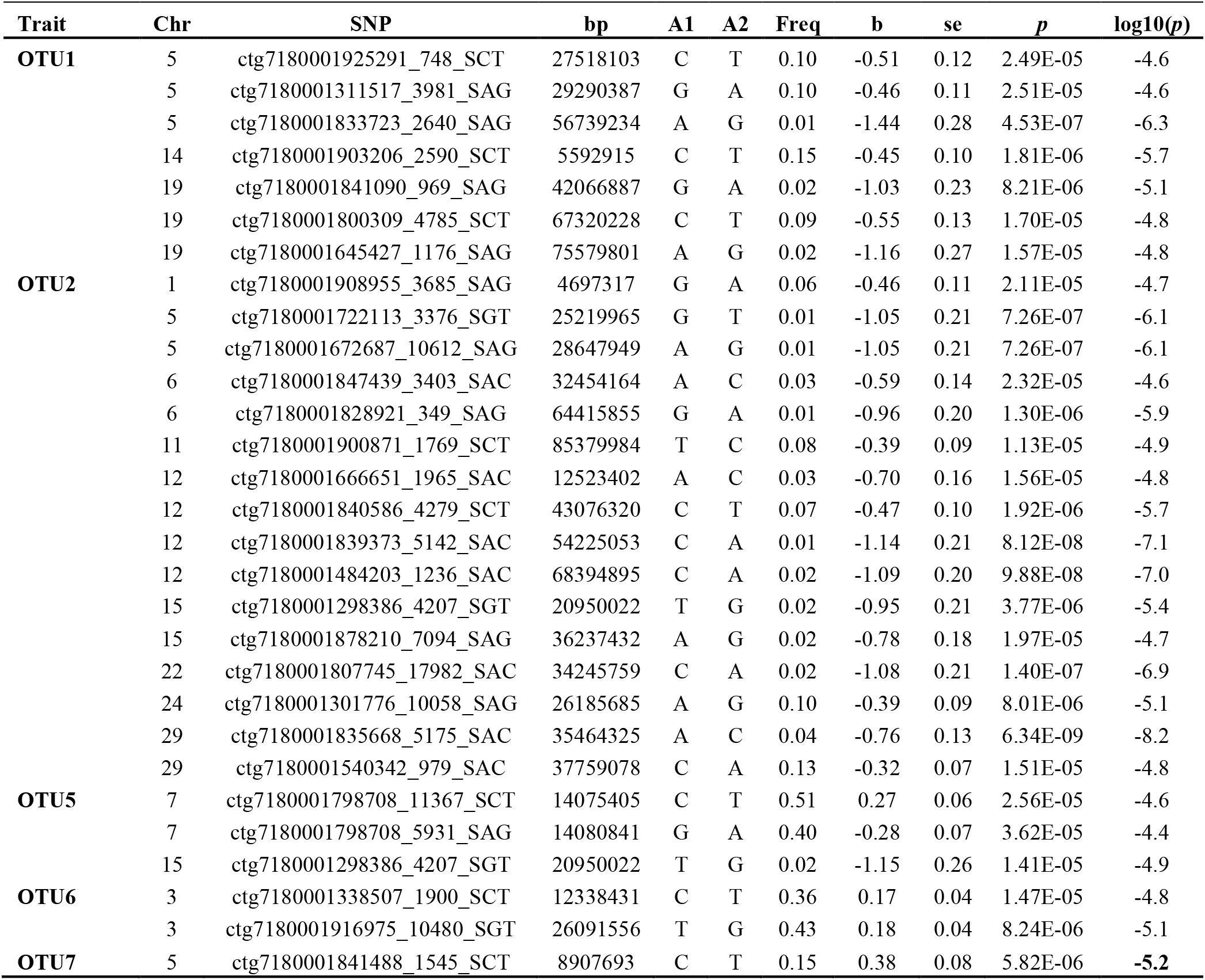
Single-nucleotide polymorphisms (SNP) associated with the OTU variables.

**Supplementary Table 2.**
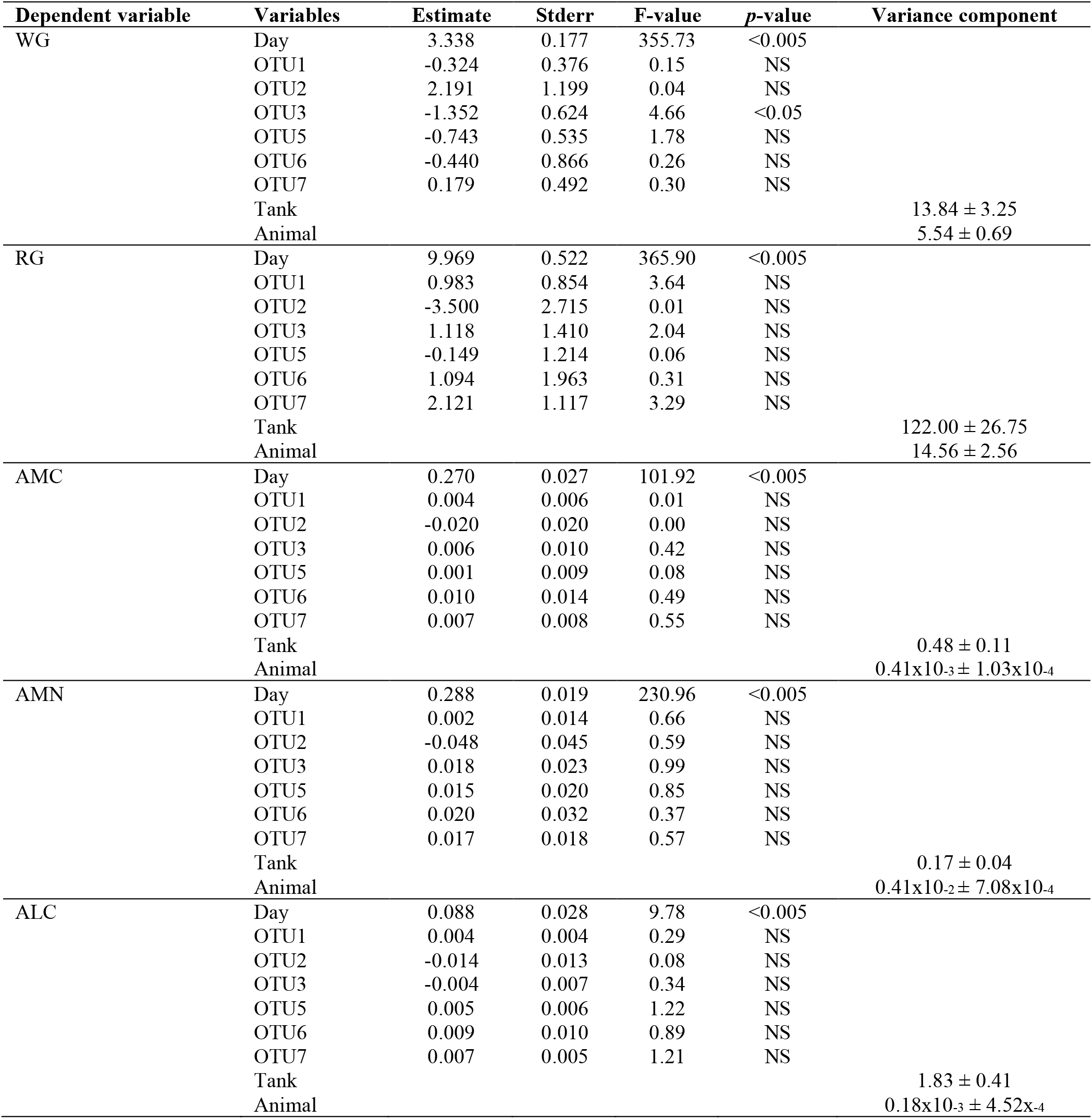
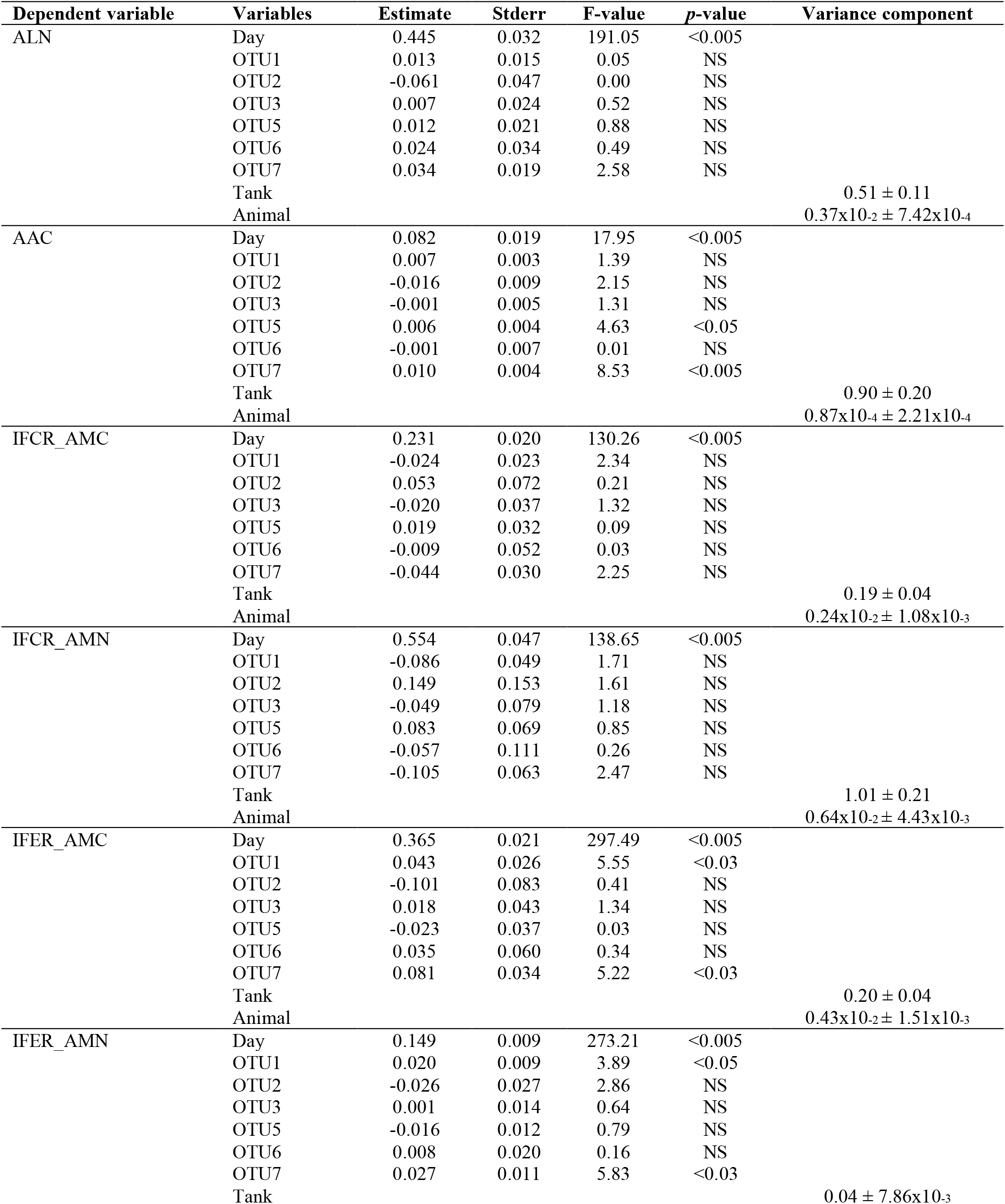
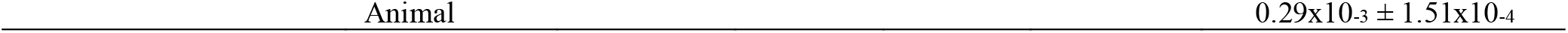
Regression estimates, standard errors, *F*- and *p*-values when regressing OTUs on growth, metabolism, and feed efficiency variables. The model also contained regression on day and random effects of animal (utilizing genomic relationships), and tank for which variance components are included.

## SUPPLEMENTARY FIGURES

**Suppl. Fig 1.**
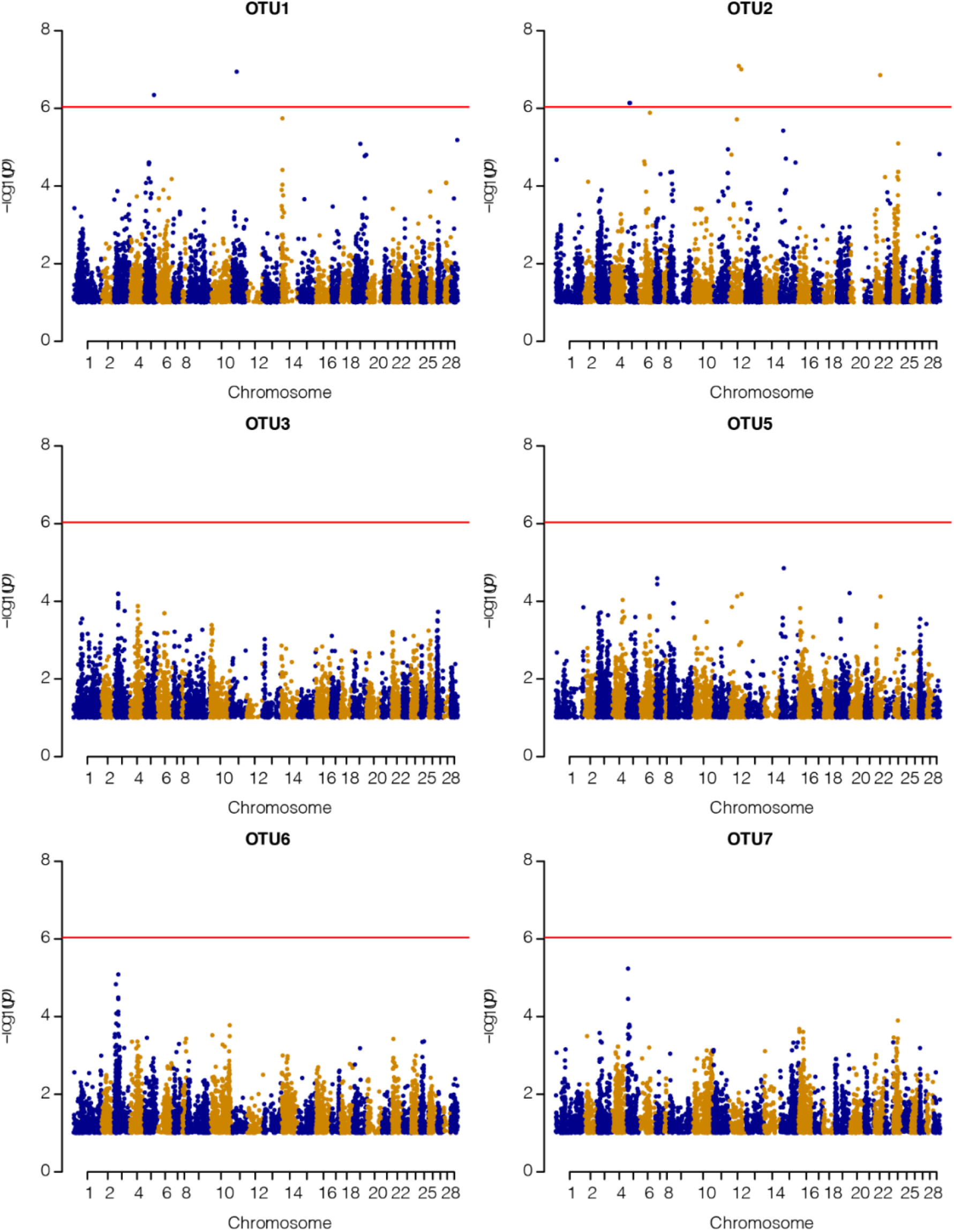
Genome-wide Manhattan plot for the different OTUs. The horizontal line represents the genome-wide Bonferroni -log_10_ (*p*) = 6.03 threshold.

**Suppl. Fig 2.**
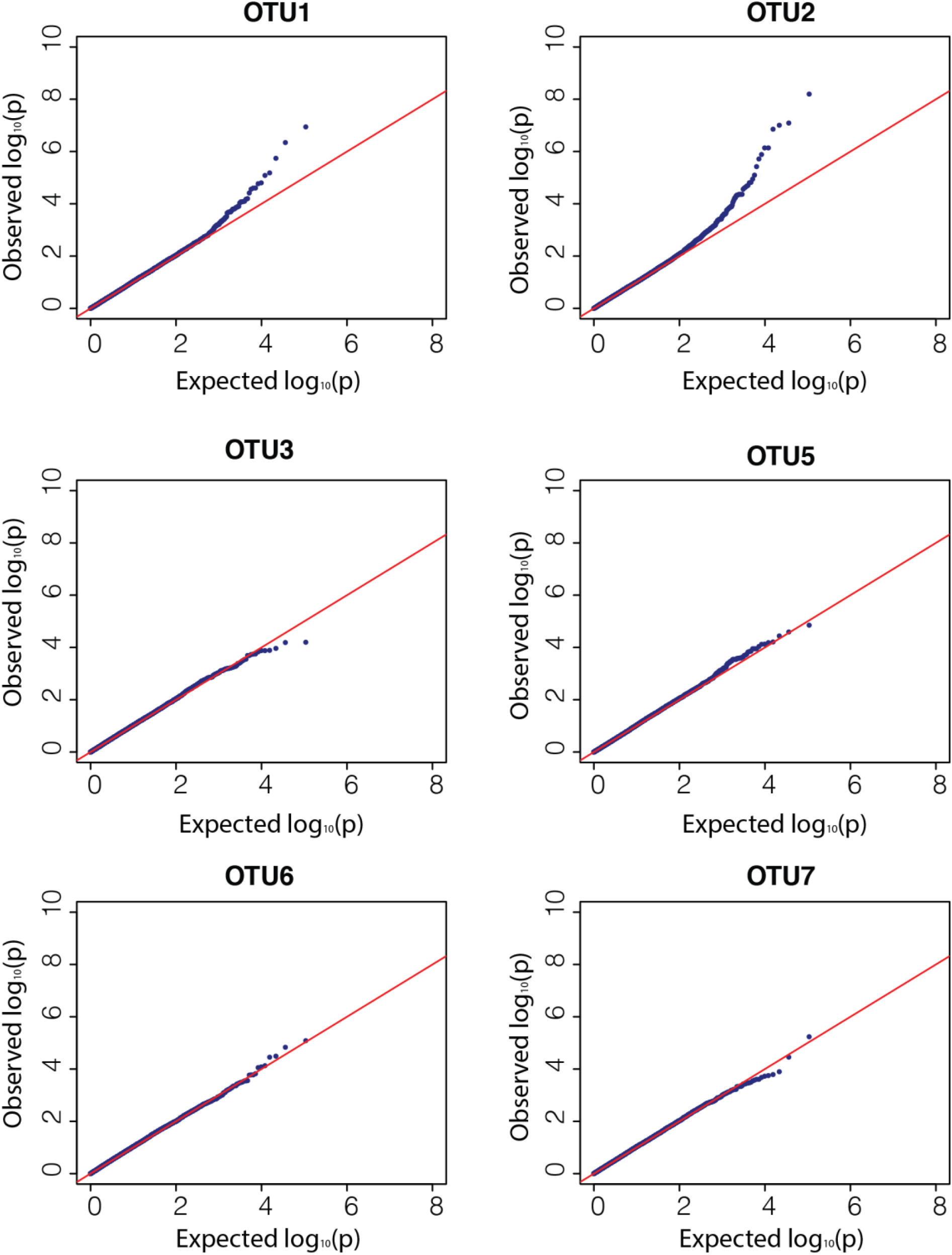
Q-Q plots from genome-wide association analyses of the different OTUs.

